# Scalable learning of interpretable rules for the dynamic microbiome domain

**DOI:** 10.1101/2020.06.25.172270

**Authors:** Venkata Suhas Maringanti, Vanni Bucci, Georg K. Gerber

## Abstract

The microbiome, which is inherently dynamic, plays essential roles in human physiology and its disruption has been implicated in numerous human diseases. Linking dynamic changes in the microbiome to the status of the human host is an important problem, which is complicated by limitations and complexities of the data. Model interpretability is key in the microbiome field, as practitioners seek to derive testable biological hypotheses from data or develop diagnostic tests that can be understood by clinicians. Interpretable structure must take into account domain-specific information key to biologists and clinicians including evolutionary relationships (phylogeny) and dynamic behavior of the microbiome. A Bayesian model was previously developed in the field, which uses Markov Chain Monte Carlo inference to learn human interpretable rules for classifying the status of the human host based on microbiome time-series data, but that approach is not scalable to increasingly large microbiome datasets being produced. We present a new fully-differentiable model that also learns human-interpretable rules for the same classification task, but in an end-to-end gradient-descent based framework. We validate the performance of our model on human microbiome data sets and demonstrate our approach has similar predictive performance to the fully Bayesian method, while running orders-of-magnitude faster and moreover learning a larger set of rules, thus providing additional biological insight into the effects of diet and environment on the microbiome.

## 1. Introduction

The human microbiome refers to the collection of trillions of micro-organisms living on and within us. The microbiome is inherently dynamic, changing due to factors including maturation of the gut in childhood, diet, environmental exposures, and medical interventions (Gerber, 2014). Recent studies have investigated the microbiome longitudinally in the context of various human diseases (e.g., diabetes (Kostic et al., 2015) or mortality after hematopoietic-cell transplantation (Peled et al., 2020)). Analyzing such time-series data is key to robustly linking the microbiome to disease and uncovering potentially causative relationships.

However, extracting meaningful information from these data sets is challenging. Human microbiome time-series studies have relatively few subjects, exhibit high degrees of subject-to-subject variability, and are typically imbalanced and irregularly temporally sampled (Gerber, 2014). Inherent aspects of microbiome data present additional challenges, including high-dimensionality, compositionality, zero inflation and complex dependencies among variables (Tsilimigras & Fodor, 2016).

Model interpretability is also very important in microbiome studies, because researchers usually seek to generate testable biological hypotheses or inform clinical decisions. A key component of interpretability in this domain is phylogeny, or information about the evolutionary relationships among microbes, which is typically encoded in a tree structure with extant microbes or Operational Taxonomic Units [OTUs] forming the leaves of the tree. Such information is used by practitioners to interpret the relatedness of microbes identified through sequencing data, which allows one to make educated guesses about the phenotypic properties of the microbes (e.g., their potential to produce immune-modulating compounds) and to define conditions for isolating the microbes via culture-based techniques. Moreover, for microbiome data that is dynamic, meaningful interpretation must also take into account temporal information, e.g., *when* in the first year of life do we see changes in the microbiome that differentiate breast-fed versus formula-fed infants?

A state-of-the-art method in the microbiome field that addresses the above-mentioned challenges is MITRE (Bogart et al., 2019), which uses a fully Bayesian model that learns human-interpretable rules to classify the host’s status (e.g., healthy or diseased) based on microbiome time-series data. MITRE rules consist of conjunctions of *detectors* that handle dependencies in both microbiome and time-series data. These detectors are of the form: *“TRUE if the aggregated abundance of organisms in phylogenetic subtree A within time window T is above threshold B*.*”* The rules are combined via logistic regression to model the probability of the classification label. This approach, which performs a set of nonlinear but interpretable and domain-specific transformations on the covariates, was demonstrated to perform generally on par or often outperform “black-box” machine learning methods (e.g., Random Forests) and produce biologically meaningful results on validation data sets To perform inference, MITRE uses a Markov Chain Monte Carlo algorithm with custom Metropolis Hastings moves that operate combinatorially on a large space of precomputed features (e.g., abundance thresholds and phylogenetic subtrees.) Consequently, MITRE’s runtime can be very slow on even moderately sized data sets, in some cases exceeding a day.

To address the challenge of scalability to large data sets while maintaining interpretability, we developed a novel 5-layer Neural Inductive Logic Programming (ILP)-type model with domain-specific microbiome and temporal attention mechanisms that outputs human-interpretable classification rules. Our method is inspired by MITRE and can be viewed as a fully-differentiable relaxation amenable to fast gradient descent-based inference. Several relaxation techniques are employed including the concrete distribution for binary selector variables (Maddison et al., 2016) to encode model sparsity and approximation of the Heaviside function to encode domain-specific attention mechanisms. We use an efficient gradient descent-based inference approach, and demonstrate on three human microbiome data sets that our model’s classification performance is similar to that of MITRE, while maintaining interpretability and running orders of magnitude faster. Moreover, our model learns a richer set of classification rules that suggest further biological insights into the effects of diet and environment on the microbiome.

## 2. Related work

### Machine learning for microbiome time-series data

Although a variety of machine learning approaches have been applied to static microbiome data (Vangay et al., 2019; Haran et al., 2019; Knights et al., 2011a;b), there remains a dearth of microbiome time-series specific methods. Typical analysis methods in the literature include performing statistical tests for differential abundance at each time-point individually or over manually-selected temporal windows of interest for each taxonomic level to find differences between two cohorts (Arrieta et al., 2015). Such approaches obviously do not effectively use temporal information and moreover cannot find interactions between microbes. Mixed-effects models have been developed for analyzing microbiome data with repeated measurements (Chen & Li, 2016; Bokulich et al., 2018). However, these approaches do not model interactions or other nonlinearities between microbes. As noted, phylogenetic structure is important for microbiome data analysis and various models have been developed that take into account phylogeny (e.g., (Tang et al., 2017)), but these methods are generally not applicable to time-series data. Our work is directly inspired by MITRE (Bogart et al., 2019), which is currently the state-of-the-art for microbiome time-series specific classification methods. However, MITRE’s MCMC-based inference algorithm suffers from scalability issues, effectively sampling over a discrete combinatoric space of phylogenetic subtrees, time windows and abundance thresholds that must be pre-enumerated. In contrast, our approach relaxes the problem to a continuous space, thus enabling use of highly efficient optimization tools.

### Differentiable rule learning

Our approach is related to recent work that seeks to unify Inductive Logic Programming (ILP) with deep learning architectures amenable to gradient-based optimization, including (Evans & Grefenstette, 2018; Yang et al., 2017; Madsen & Johansen, 2020). ILP is a broad field concerned with learning logical rules from data, often with a goal of inferring human interpretable representations. For example, Differentiable Inductive Logic Programming (Evans & Grefenstette, 2018) proposes a framework to learn symbolic rules on small noisy data sets and can also generalize to non-symbolic domains such as images. Other examples include Neural Logic Programming (Yang et al., 2017), a framework that learns first-order logical rules for mining knowledge bases and Neural Arithmetic Units (Madsen & Johansen, 2020) that learns basic arithmetic operations that can extrapolate well to numbers out of the range of training data. These methods share a common theme with our model in that they induce direct and task-specific inductive biases with the goal of improving generalization, robustness and interpretability. In our case, we incorporate into our model inductive biases via temporal and phylogenetic attention mechanisms that leverage the domain-specific structure of microbiome time-series data.

### Interpretability in deep models

Deep learning models have achieved state-of-the-art predictive performance on many tasks in fields such as computer vision, natural language and speech processing. Although predictive performance is an important metric, model interpretability is also essential as these systems are increasingly being deployed in high-risk scenarios for augmenting human decision making (Lipton, 2016; Doshi-Velez & Kim, 2017). A variety of *post hoc* interpretation methods or methods that remove features and evaluate the impact on the model have been described, but the effectiveness of these methods remains unclear, e.g., (Hooker et al., 2019). Our work differs from these methods by constructing a model encoding a specific structure that is inherently interpretable throughout the layers and in the final output.

## 3. Model

### 3.1. Overview

Our model classifies hosts (e.g., individual human subjects) given temporal, high-dimensional measurements of their microbes, while incorporating domain-specific inductive biases to facilitate interpretability. This structure includes aggregation of covariates into features that preserve phylogenetic and temporal information in the data and selection of a sparse set of features. Our model learns which covariates to attend to for the classification task, with attention influenced by the dependence structure among the covariates.

Figure 1A shows our model as a graphical (plate) representation and Figure 1B shows it in layered form. Let *Y*_*s*_ denote the binary label for subject *s*. Let *X*_*sit*_ denote a microbiome measurement from subject *s* at time *t* for OTU *i* of an *N* -dimensional data source (e.g., relative abundances of OTUs from 16S rRNA amplicon or metagenomic shotgun sequencing). The task is then to output the probability of the label for each subject given the data for that subject, e.g., *P* (*Y*_*s*_ | **X**_*s*_). We further assume there exists a graph structure on the OTUs, such that *d*_*ij*_ specifies the weight of the edge between OTU *i* and *j* (i.e., derived from a phylogenetic tree.)

**Figure 1.**
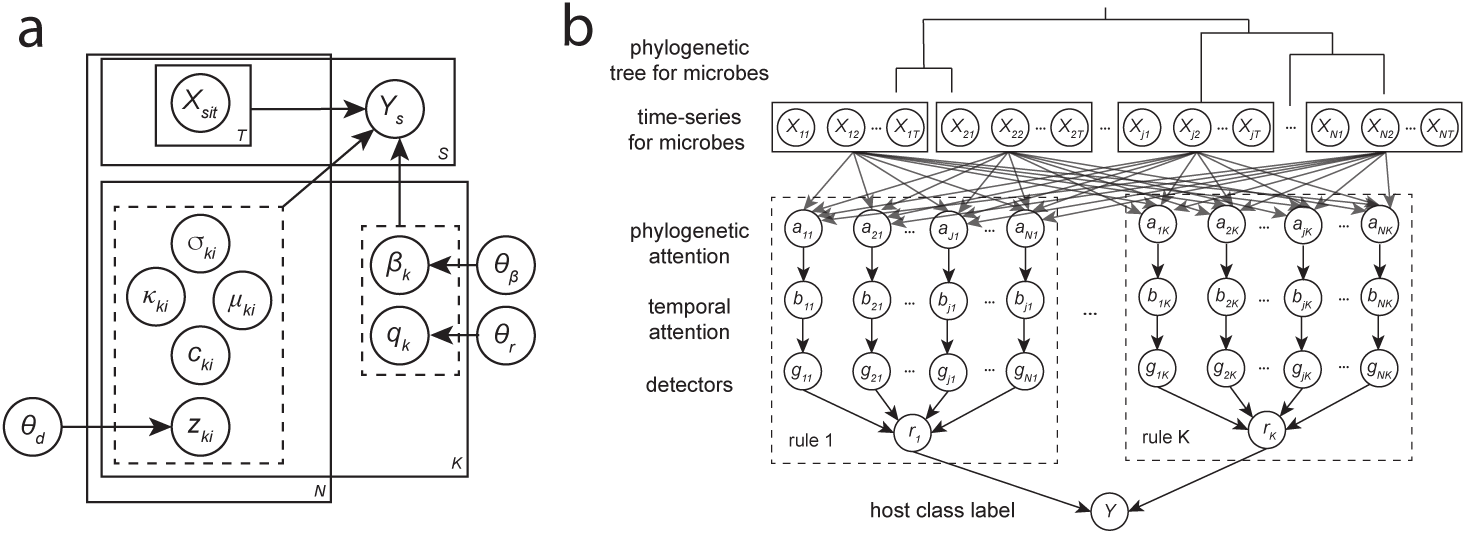
**(a) Our probabilistic model**. The binary class label *Y*_*s*_ for each subject is dependent on the input data *X*_*sit*_ (microbiome time-series) and variables for *K* rules, each comprised of variables for *N* detectors. Indicator variables *q* and *z* select which rules or detectors are active and have priors with parameters *θ*_*r*_ and *θ*_*d*_ respectively. Rule weights *β* have a prior with parameters *θ*_*β*_. Detector variables *µ* and *σ* specify temporal attention filter centers and bandwidths respectively; *κ* specify phylogenetic attention filter bandwidths; and *c* specify detector thresholds. **(b) Layered representation of our model**. Our model can be viewed as a 5-layer Neural ILP-type model that assigns labels to hosts given input microbial time-series data and a graph structure based on phylogenetic relationships between bacteria. The outputs of each layer are composed to form the final output of human-interpretable rules that produce the classification label.

We model *P* (*Y*_*s*_ | **X**_*s*_) as a logistic regression over rules. Each rule is comprised of conjunctions of *detectors* of the form *“(SOFT) TRUE If the aggregated abundance of organisms in phylogenetic cluster A within time region T is above threshold B*.*”* Note that in the original MITRE model that served as an inspiration for our model, strict phylogenetic subtrees are used, whereas in our model we use clusters (e.g., nearby OTUs in phylogenetic space) to facilitate differentiability. Our model additionally relaxes strict time windows to time regions and hard thresholds to soft decision boundaries. Associated with each relaxation is a sharpness parameter. These parameters are annealed during our gradient descent-based learning algorithm, to arrive at detectors that approximate the discrete nature of the original MITRE detectors. We implemented our model in PyTorch (Paszke et al., 2019); source code is available at https://github.com/gerberlab/mditre.

### 3.2. Rules

We assume a maximum of *K* rules with each rule comprised of a maximum of *N* detectors. Let *g*_*ki*_(·) denote binary valued detector *i* in rule *k* and *z*_*ki*_ denote a binary variable that selects whether the detector is active (value of 1) in the rule. A binary valued *hard* rule *r*_*k*_(·) would be the logical conjunction of active detectors:

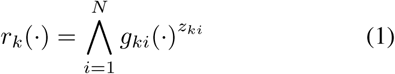

In order to make our model differentiable, we relax the logical conjunction using the following approximation:

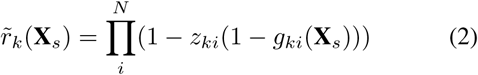

This formulation is inspired by the Neural Multiplication Unit from (Madsen & Johansen, 2020). Additionally, we relax *z*_*ki*_ by modeling it as a binary concrete (Maddison et al., 2016) variable.

As described below, the detector functions *g*_*ki*_(·) will effectively learn focused features from the data: (a) localizing regions in the phylogenetic space and, (b) time-periods in the time-series.

### 3.3. Detectors

#### 3.3.1. Phylogenetic Attention Mechanism

We use a relatively sharp (but smooth) averaging filter that serves as an attention mechanism over the graph on OTUs:

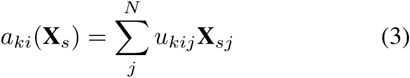

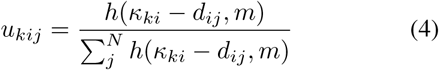

Here, *u*_*kij*_ is a normalized weight for OTU *j* and *κ*_*ki*_ controls the filter bandwidth. The function *h* is an analytic approximation to the Heaviside step function:

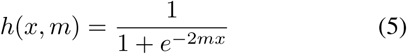

The annealing parameter *m* controls the sharpness of the response, i.e., in the limit of large *m* the detector will include all OTUs within a radius *κ*. Intuitively, this filter “pulls in” other OTUs phylogenetically close to central OTU *i*.

#### 3.3.2. Temporal Attention Mechanism

We similarly use a relatively sharp (but smooth) averaging filter which serves as an attention mechanism over time:

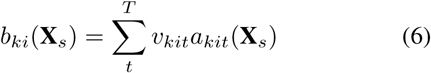

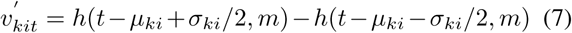

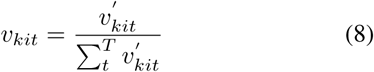

Here, *µ*_*ki*_ controls the center of the filter and *σ*_*ki*_ its bandwidth. As the annealing parameter *m* increases, this will approach a time window of duration *σ* and center *µ*.

#### 3.3.3. Thresholds

We model the response of detector *g*_*ki*_ as:

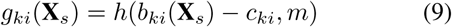

Here, *c*_*ki*_ specifies the value for a soft-threshold. Again, as the annealing parameter *m* increases, this will approach a hard threshold.

### 3.4. Classifier

Finally, we model the class probability for subject *s* via logistic regression with the rules as covariates:

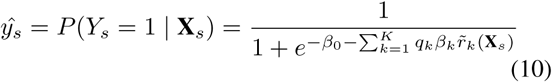

Here *q*_*k*_ is a binary concrete (Maddison et al., 2016) variable that selects which rules to include in the model.

### 3.5. Priors and Loss Function

To encourage parsimonious classifiers, we place priors on the number of rules and detectors. Specifically. we place Negative Binomial Distributed (NBD) priors on the total number of active rules 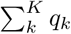 and the number of active detectors per rule 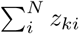 with mean 0 and variance 100 in both cases, thus encoding sparsity with a prior belief of no rules or detectors. We also place a diffuse Gaussian prior on the regression coefficients *β*_*k*_ with mean 0 and variance 10^5^; for other parameters we assume improper flat priors.

Our loss function is then equal to the negative log likelihood of the posterior and we perform maximum a posteriori (MAP) estimation:

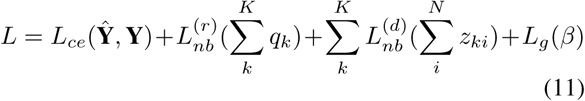

Here, *L*_*ce*_ is the standard cross-entropy loss, 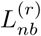 and 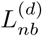 are the negative logs of the NBD priors and *L*_*g*_ is the negative log of the Gaussian prior.

## 4. Experiments

### 4.1. Data sets

We evaluated the performance of our model on human microbiome data sets originally assessed by MITRE (Bogart et al., 2019), using the same preprocessing and class labels as in the original study, in order to provide comparable results. These data sets consist of 16S rRNA amplicon sequencing data, which identifies the relative abundances of OTUs in the sample. **David** et al. (David et al., 2014) (20 subjects, 185 OTUs, class label = plant vs. animal diet) studied the effects of short-term dietary changes on the human gut microbiome. They tracked the gut microbiome composition of healthy adults before, during, and after a 5-day period of consuming exclusively plant-based or exclusively animal-based diets. **Bokulich** et al. (Bokulich et al., 2016) (35 subjects, 60 OTUs, class label = formula vs. breastfed diet) studied the effects of antibiotic exposures, cesarean section, and formula feeding on the gut microbiome composition during the first 2 years of infancy. **Vatanen** et al. (Vatanen et al., 2016) (113 subjects, 237 OTUs, class label = Russian vs. Estonian/Finnish nationality) tracked the gut microbiome composition from birth to 3 years of age in cohorts of children at high risk for autoimmune disease.

### 4.2. Model training details

We set *K* = 10 and *N* = # OTUs for the maximum number of rules and detectors per rule respectively. We used full-batch gradient descent with the Adam optimizer (Kingma & Ba, 2014) and cosine learning rate annealing schedule over 1000 training epochs. We used a nested 5-fold cross-validation scheme to determine the number of epochs to train. The sharpness parameter *m* of the Heaviside step function approximation and temperature of the binary concrete variables are linearly annealed during training. Our model was run on a single Tesla V100 GPU with 32-bit precision training. Please see our source code for full details on training, including the initialization used.

### 4.3. Performance Results

We used cross-validated F1 scores (harmonic mean of precision and recall), as in the original MITRE paper, to assess generalization performance. Table 1 shows the performance and runtime comparison between MITRE and our model. Our results indicate that our differentiable model performs quite similarly to the fully Bayesian method, while running approximately 30-70X faster, suggesting our approach provides greater scalability without sacrificing performance.

**Table 1.** Cross-validated F1 score and runtime comparison for MITRE and our model (DR = differentiable rules) on the data sets analyzed. We report the mean and standard deviation of F1 score for our model on 10 random seeds, with the runtime given in parenthesis. Scores and runtime of MITRE are taken directly from (Bogart et al., 2019), which provided only mean performance statistics.

| MODEL | DAVID<br>(DAVID ET AL., 2014) | VATANEN<br>(VATANEN ET AL., 2016) | BOKULICH<br>(BOKULICH ET AL., 2016) |
| --- | --- | --- | --- |
| MITRE | 0.77 [23 HRS] | 0.83 [64.50 HRS] | 0.64 [30 HRS] |
| DR | 0.81 ± 0.07 [0.45 HRS] | 0.82 ± 0.05 [0.83 HRS] | 0.58 ± 0.1 [1 HR] |

### 4.4. Model Interpretability

We show that our model learns rules that are human-interpretable and moreover provide insights in the microbiome domain not seen with the fully Bayesian method. We note also that our model learned a different number of rules and average number of detectors per rule on each of the data sets (see below), suggesting that it automatically expands model capacity as needed. This capability is part of the inference procedure and does not require seperate hyperparameter tuning sweeps, which would be very expensive over the large combinatorial space.

**David** (David et al., 2014). Our model learned 2 rules with 1 detector per rule. Both detectors are temporally meaningful: the learned time windows occur during the dietary intervention (day 5 to day 10). Moreover, the detectors are biologically meaningful and provide phylogenetic context that greatly facilitates interpretation. The first rule/detector reads: *“TRUE if the aggregated relative abundance of OTUs [Operational Taxonomic Units] in the family Eubacteriacaeae and in the genera Roseburia is above 0*.*49% between day 7 and day 10”* and is similar to one found by MITRE. The second rule/detector (Figure 2a) reads: *“TRUE if the aggregated relative abundance of OTUs from the species Bilophila wadsworthia and genera Desulfovibrio is above 0*.*31% between day 7 and 11*.*”* This detector was not found by MITRE, but is clearly biologically meaningful. The detector differentiates subjects on the animal-based diet by increased abundances of two taxa belonging to the species *Bilophila wadsworthia* and to the genus *Desulfovibrio. B. wadsworthia* is a sulfite-reducing pathobiont shown to increase in abundance with dietary lipids (Devkota et al., 2012) and *Desulfovibrio* is a sulfate-reducing bacterium shown to increase in abundance with chondroitin sulfate (an important component of animal cartilage) (Rey et al., 2013).

**Figure 2.**
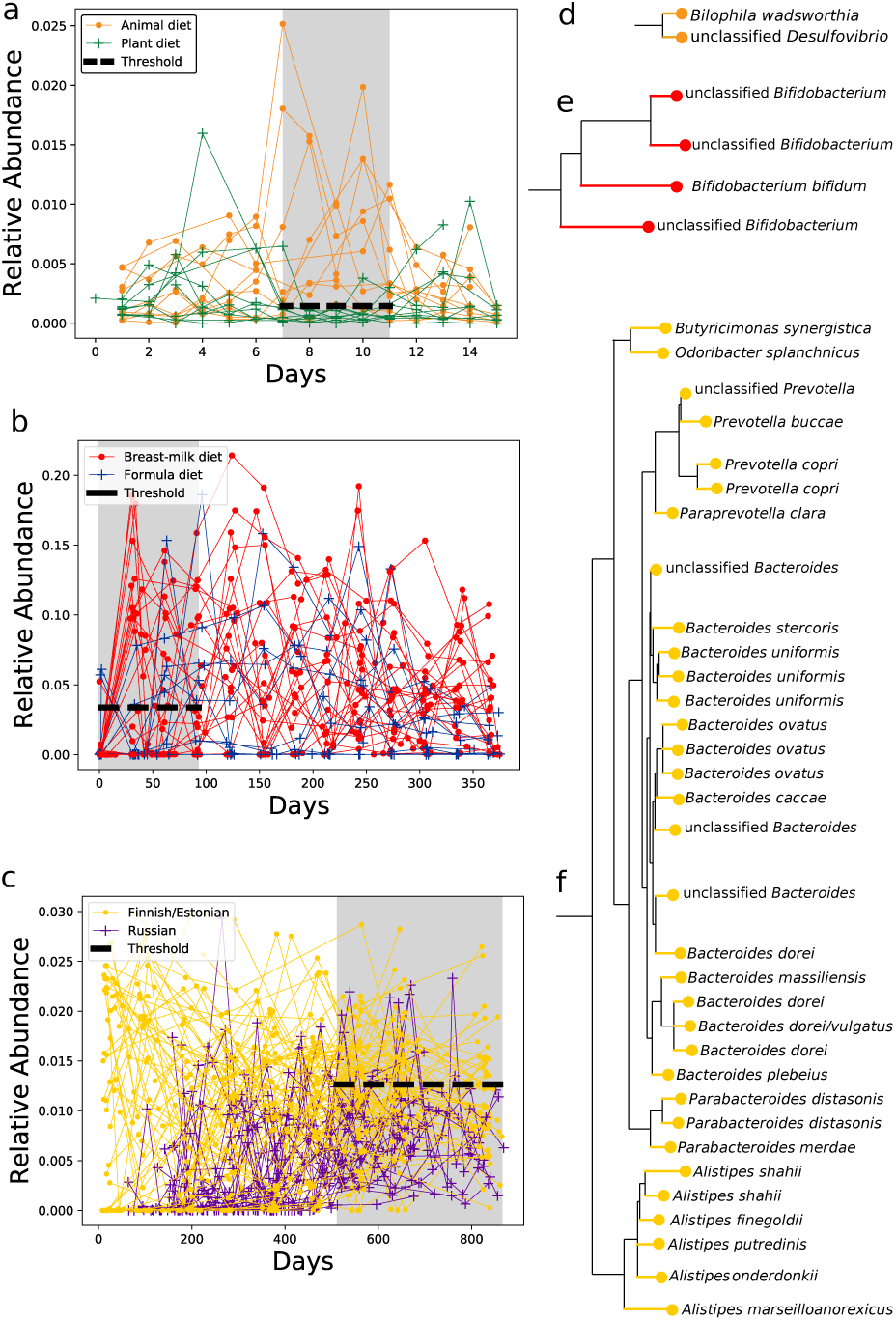
Visualization of example detectors. Panels (a-c) show detector time-windows (shaded region) and thresholds (black dashed line) superimposed on data (aggregated relative abundances over time for Operational Taxonomic Units [OTUs] selected by the detector.) Panels (e-f) show the OTUs selected by the detector and their phylogenetic context. Figures (a) and (d): detector discriminating study participants eating plant versus meat-based diets, (b) and (e): detector discriminating infants on breast-milk versus formula predominant diets, (c) and (f): detector discriminating children of Russian versus Finnish/Estonian nationalities.

**Bokulich** (Bokulich et al., 2016). Our model learned 5 rules with an average of 1.5 detectors per rule. In the interest of space, we discuss one of the single detector rules that is clearly biologically relevant and was not found by MITRE. This rule is true mostly for infants on a predominantly breast-milk diet and reads, *“TRUE if the aggregated relative abundance of OTUs in the genus Bifidobacterium is above 3% between day 0 and day 93*.*”* This rule is clearly biologically meaningful, as bacteria in the genus *Bifidobacterium* have been shown to express the necessary enzymes to digest human milk oligosaccharides (Lawson et al., 2020).

**Vatenen** (Vatanen et al., 2016). Our model learned 3 rules with 1 detector per rule. The first rule/detector that reads, *“TRUE if the aggregated relative abundance of OTUs from the genera Lactobacillus, Streptococcus, Granulicatella, Enterococcus and Staphylococcus is above 0*.*17% between day 208 and day 466”* is similar to one found by MITRE. The second two rules/detectors were distinct from ones learned by MITRE. These rules/detectors, which are true predominantly for subjects of Russian nationality read *“TRUE if the aggregated relative abundance of OTUs from the genera Alistipes, Bacteroides, Butyricimonas, Parabacteroides, Prevotella is below 0*.*9% between day 385 and day 659”* and *“TRUE if the aggregated relative abundance of OTUs from the genera Alistipes, Bacteroides, Butyricimonas, Parabacteroides, Prevotella is below 1*.*2% between day 511 and day 886*.*”* These rules indicate relatively depleted levels of *Bacteroides* and related bacteria in the second year of life, which may reflect different degrees of plant fiber consumption between children in Russia vs. Finland/Estonia after switching to exclusively solid foods (Vatanen et al., 2016).

## 5. Conclusion

We have presented a fully differentiable model with domain-specific microbiome/temporal inductive biases that learns classifier rules and achieves competitive generalization performance with a fully Bayesian model, MITRE, while running 30-70X faster. We have further demonstrated that our method learns human-interpretable rules that are biologically meaningful, including rules not found by MITRE. A possible reason that our model learns a richer set of rules is that MITRE effectively treats the problem combinatorially, i.e., sampling from different discrete thresholds, phylogenetic subtrees, etc. In contrast, our model relaxes the problem and has a much faster run-time, which appears to allow broader exploration of the space of rules, while still learning sharp/almost discrete structure. Of note, our model automatically expands capacity (number of detectors and rules) as part of the core inference procedure, without requiring exogenous and expensive hyperparameter tuning. Directions for future work include specifying additional priors in the model, for instance to encourage time windows or phylogenetic aggregation based on prior beliefs of biologist colleagues. Another direction would be to incorporate different types of detectors, including rate-of-change of abundance (as in the original MITRE model) or types of detectors tailored to new datatypes such as metabolomic or metatranscriptomic information. Also, although our MAP inference algorithm performed well, it would be interesting to explore Variational Inference to obtain approximations to the model posterior. Finally, we intend to perform experiments on additional microbiome data sets that are already published as well as new data that are currently being collected from large cohorts of hospitalized patients.

